# A large pocket structure surrounding the catalytic center in the BCG protein from *Mycobacterium tuberculosis*

**DOI:** 10.1101/2024.05.14.591795

**Authors:** Kohei Takeshita, Naoki Sakai, Go Ueno, Midori Horie, Hirofumi Tsujino, Mitsuhiro Arisawa, Masaki Yamamoto, Masayoshi Arai, Taku Yamashita

## Abstract

The widespread prevalence of tuberculosis and the need for extensive chemotherapy in its treatment are caused by the ability of tuberculosis (TB) bacteria to enter a dormant state. Therefore, it is crucial to develop effective compounds that can inhibit the growth of TB bacteria in both the active and dormant states. Previous research has identified agelasine D (marine sponge-derived diterpene alkaloid) capable of resisting dormant mycobacteria. Additionally, BCG3185c has been designated as a target mycobacterial protein. In this study, the crystal structure of BCG3185c was determined at a resolution of 1.86 Å. Analysis of this crystal structure revealed a large pocket toward the catalytic center of BCG3185c, which is potentially adequate for agelasine D binding. Based on the results of the interaction analysis between agelasine D and *Bacillus* Calmette-Guérin (BCG) proteins, it was inferred that the binding of agelasine D to this large pocket is crucial for suppressing the growth of *Mycobacterium tuberculosis*.

## Introduction

Tuberculosis (TB) is caused by *Mycobacterium tuberculosis*, and it spreads when an infected person releases the bacteria into the air. It is estimated that approximately a quarter of the world’s population is infected with TB. Although preventable and usually treatable, as of 2022, it ranks as the second leading cause of death worldwide from a single infectious pathogen, following the novel coronavirus infection (COVID-19), resulting in almost twice as many deaths as human immunodeficiency virus/acquired immunodeficiency syndrome (HIV/AIDS) ^1^.

Additionally, factors, such as fragile healthcare system environment, emergence of multidrug-resistant strains of tuberculosis, and dual infection with HIV/AIDS or emerging resurgence viruses have escalated TB into a significant health concern. TB is one of the leading causes of death, specifically in areas where HIV infection is endemic ^2^. *Mycobacterium tuberculosis* exhibits a challenging characteristic of adapting to the environment at the site of infection, called granuloma, and entering a dormant state, thereby requiring prolonged treatment (a minimum of six months). Therefore, it is desirable to develop novel lead compounds with growth-inhibitory activity against both the proliferative and dormant states of *Mycobacterium tuberculosis*. Because hypoxia can induce the dormant state of *Mycobacterium tuberculosis* ^3-5^, Arai et al. developed a screening method to identify antimicrobial substances that are effective against the dormant state of *Mycobacterium* spp., and successfully isolated active substances, such as halicyclamines (macrocyclic alkaloids) from *Haliclona* sponge ^6,7^, sponge-derived *Trichoderma* fungal cultures, trichoderin (novel aminolipopeptide) ^8^, and neamphamide B (novel cyclic depsipeptide) ^9^ from *Neamphius* sponges. Additionally, the screening of compounds from marine organisms revealed agelasines B, C, and D ^10,11^ from the Indonesian marine sponge (genus *Agelas*), which are effective against dormant mycobacteria. Moreover, agelasines characterized by diterpene alkaloid structures exhibit various activities, such as antibacterial ^12-17^, antiproliferative ^15-18^, antiprotozoal ^19,20^, anti-biofilm formation ^16^, anti-staining ^16^, and Na+/K+-ATPase inhibition in mammalian tissues ^10,21^. Specifically, agelasine D exhibited antimicrobial activity against *Mycobacterium tuberculosis* H37Rv during active growth, and BCG3185c was identified as its target molecule in 2014 ^22^. BCG3185c is anticipated to be part of the Rieske-type dioxygenase family because amino acid sequence analysis demonstrated that it contains a Rieske iron-sulfur cluster domain and is homologous to the α subunit of dioxygenase. This family of enzymes is abundant in soil bacteria, such as *Pseudomonas*, and their reactions are responsible for the initial steps in the biodegradation of aromatic hydrocarbons. However, BCG3185c is an orphan enzyme with unknown substrates, thereby posing a challenge for the development of novel compounds with anti-TB activity.

In this study, we determined the precise structure of BCG3185c and clarified the binding mechanism of agelasine D, which inhibits *Mycobacterium tuberculosis*. We aimed to contribute to the development of novel anti-TB compounds.

## Results and discussion

### Overall Structure of BCG3185c

The crystal structure of BCG3185c (apo-form) was determined using the molecular replacement method at a resolution of 1.86 Å (Figure 1a, Supplementary Table 1). Although the crystal contained two asymmetric molecules, it is reasonable to infer that BCG3185c acts as a trimer based on its elution volume through size exclusion chromatography (Figure 1b) and the trimeric structures of other Rieske-type dioxygenases. The crystals of BCG3185c belong to space group *P*2_1_3, with the central axis of the BCG3185c trimer located on the threefold axis of its cubic crystal (Figure 1c). This indicated that BCG3185c also forms a trimeric structure in solution. Additionally, each protomer of BCG3185c forms a trimer through the proximity assembly of the 2Fe-2S cluster region and metal-coordinated active center, facilitating electron transfer from the Fe-S cluster to the metal ion-coordinated active center, indicating that it is a typical Rieske-type dioxygenase. Subsequently, similar structures on the Dali server revealed that the structure of each BCG3185c trimer promoter was similar to that of naphthalene dioxygenase (PDB ID: 1o7g), which is a Rieske-type dioxygenase. Additionally, it is anticipated to transfer electrons from the Fe-S cluster to the catalytic Fe at the center through Y121 (Figure 1d). Although the natural substrate of BCG3185c is unknown, these structural data indicate that BCG3185c acts as a Rieske-type dioxygenase in *Mycobacterium tuberculosis*.

**Figure. 1.**
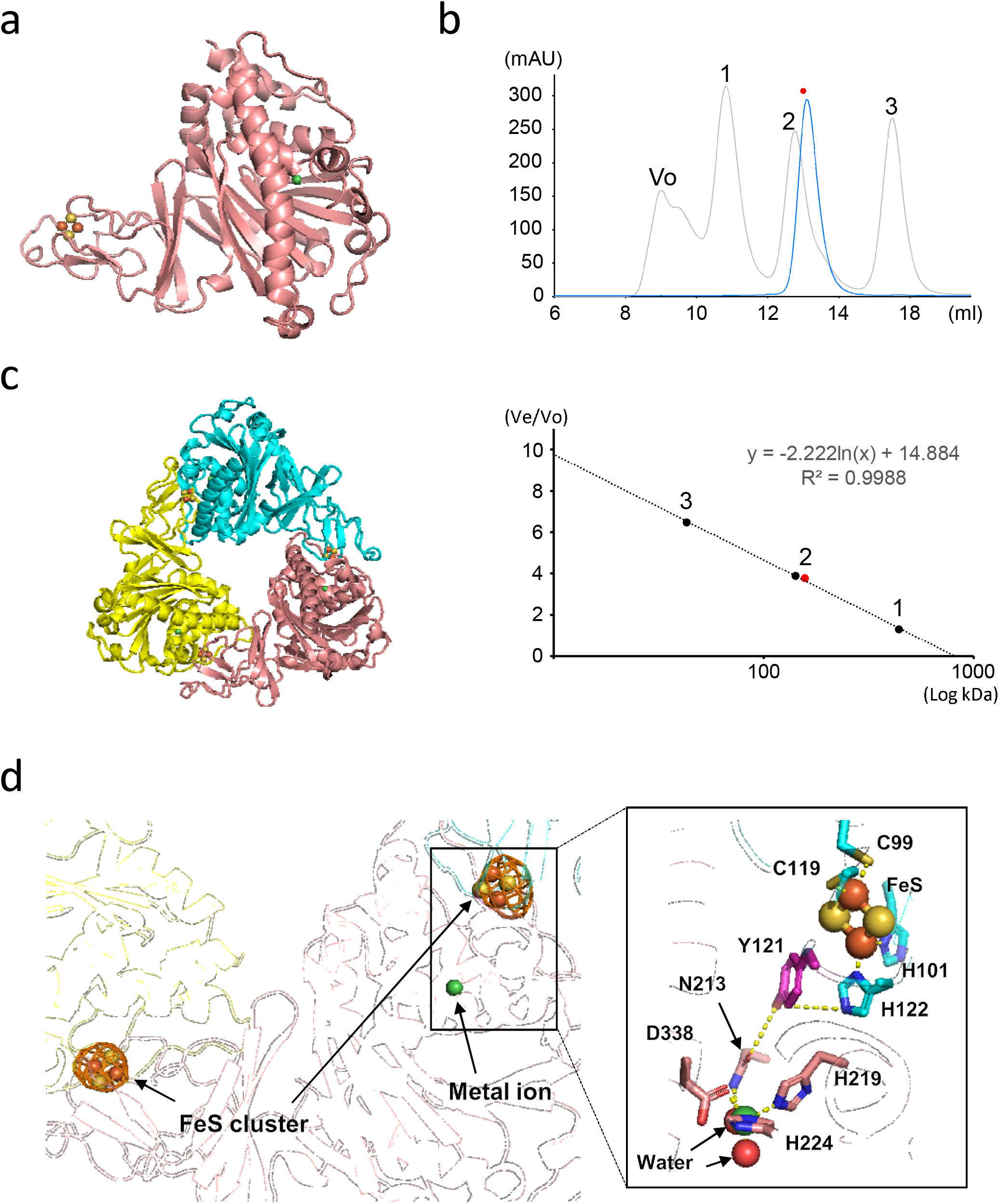
Overall structure and stoichiometry of BCG3185c in solution. (a) Crystal structure of the BCG3185c monomer. (b) Size-exclusion chromatography elution profile and standard curve of marker proteins (1: ferritin, 440 kDa; 2: aldolase, 158 kDa; 3: ovalbumin, 43 kDa; and Vo: blue dextran 2000) and BCG3185c. The standard curve is plotted with Ve/Vo on the Y-axis and the molecular weight as a logarithm on the X-axis. Vo represents the void volume and Ve represents the elution volume of each molecular weight marker protein. The elution volume of BCG3185c is indicated by an asterisk, and the Ve/Vo value is 3.87. The red circles in the size exclusion chromatography (SEC) chart and standard curve indicate the BCG3185c protein. The solution phase of BCG3185c is determined to be 142.24 kDa from the equation of the approximated standard curve (the monomer of BCG3185c has a molecular weight of 42.80 kDa). (c) Trimeric structure of BCG3185c. The threefold axis of BCG3185c is present on the threefold symmetry axis in the crystal. (d) The interaction surface between each protomer forms a structure close to the Fe-S cluster and the catalytic center of the other protomer. The Fe-S cluster is coordinated by two cysteine (C99 and C119) and two histidine (H101 and H122) residues, whereas the metal ion at the catalytic center is coordinated by two histidine residues (H219 and H224), asparagine (N213), and aspartic acid (D338) through a water molecule.

### Putative substrate binding pocket and agelasine D binding mechanism

Crystal structure analysis indicated that BCG3185c is a Rieske-type dioxygenase, and inhibition of its activity inhibits the growth of *Mycobacterium tuberculosis*. Therefore, the development of specific inhibitors based on the atomic-level structural data of BCG3185c may be crucial in the development of novel anti-TB drugs. In 2009, Arai et al. reported that agelasine D, a compound that targets BCG3185c, exerts anti-TB effects. Elucidating the binding mechanism between BCG3185c and agelasine D is crucial for the development of novel anti-TB drugs. Agelasine D is a relatively bulky natural compound with naphthalene- and purine-like structures. Because the substrates of common Rieske-type dioxygenases are low-molecular-weight compounds, such as naphthalene, it was unclear whether agelasine D may bind to BCG3185c, which is also a Rieske-type dioxygenase. Surprisingly, a large pocket was discovered that extended from the catalytic center of BCG3185c toward the surface of the molecule, which can accommodate a medium-sized natural compound, such as agelasine D (Figure 2a). This pocket is absent in the 1o7g, which has high structural similarity to BCG3185c (Figure 2b). We confirmed that there was an adequate space for agelasine D to bind to BCG3185c. Additionally, we attempted to determine the complex structure of BCG3185c and agelasine D through soaking or co-crystallization, but were unsuccessful. This may stem from the low solubility of agelasine D, which causes precipitation during the co-crystallization and soaking processes. Additionally, the crystallization conditions for BCG3185c contained high concentrations of ammonium chloride and octylphenol poly(ethylene glycol) ether (MEGA-8) surfactants. Therefore, we assessed the affinity between BCG3185c and agelasine D using microscale thermophoresis (MST) and calculated the dissociation constant (Kd) of agelasine D for BCG3185c (60 nM) to be approximately 130 μM (Figure 3a, b). The MST measurements were conducted thrice, resulting in an average Kd value (mean ± standard error of the mean [SEM]) of 150.33 ± 18.62. These results highlight the current challenges in determining the complex structure of BCG3185c and agelasine D. Because of the challenges in determining the complex structure of BCG3185c and agelasine D, docking and molecular dynamic simulations were performed to predict the binding sites of agelasine D in BCG3185c. The docking simulation yielded two potential binding sites: a large pocket (site A) connected to the catalytic center of BCG3185c and the molecular surface (site B) (Supplementary Figure 1a). At site A, the predicted binding mechanism of agelasine D was a purine-like structure bound to the catalytic center, and a naphthalene-like structure bound at the pocket entrance. Analysis of the surrounding amino acids demonstrated the presence of hydrophobic residues, indicating an interaction between agelasine D and amino acids through hydrophobic interactions (Figure 3c). Additionally, 1o7g lacks a molecular cavity similar to BCG3185c, and the substrate pocket is narrow, which may prevent agelasine D from entering. This additional feature supports the idea that agelasine D specifically binds to BCG3185c with a widened cavity. Additionally, site B was bound to the surface of the molecule. Although it is possible that the mechanism of agelasine D binding to site B and functioning as an inhibitor may disrupt the trimeric structure of BCG3185c, the binding of agelasine D to site B does not appear to inhibit its trimer formation (Supplementary Figure 1b). Therefore, it is likely that site A is the current binding site for agelasine D. Although mutant analysis can confirm this, the binding pocket is a wide hydrophobic pocket, making it challenging to detect alterations in the affinity of agelasine D without mutating most amino acids. Therefore, to clarify the binding mechanism of BCG3185c and agelasine D in the future, it is necessary to synthesize agelasine D derivatives with enhanced hydrophilicity and other variants and clarify the crystal structure of the complex with BCG3185c. These research efforts may result in the development of novel anti-TB drugs with agelasine D as the lead compound.

**Figure. 2.**
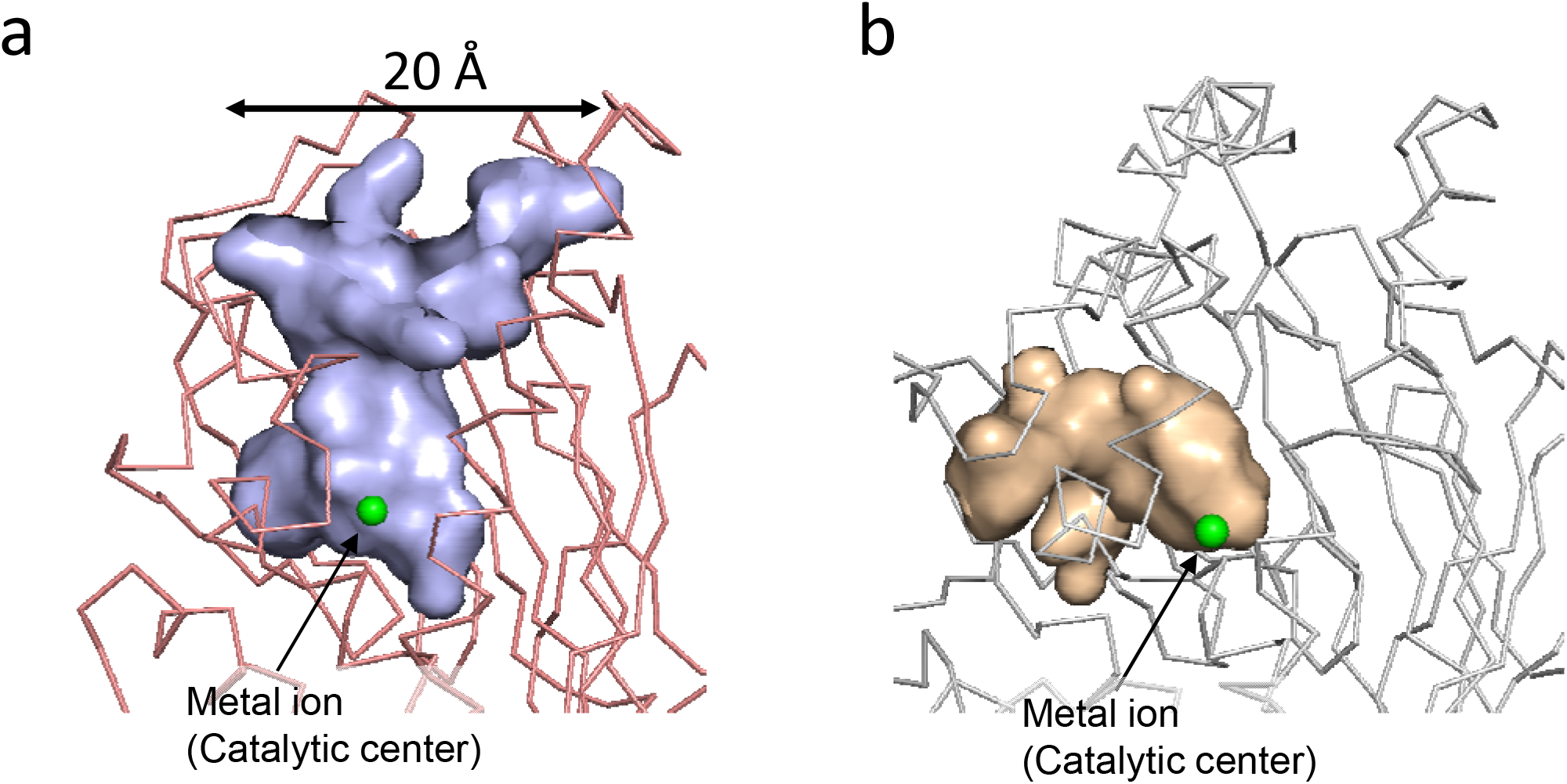
Intramolecular cavity as a putative substrate-binding pocket of BCG3185c and intramolecular cavity of naphthalene dioxygenase. (a) Intramolecular cavity observed in BCG3185c. The cavity has an entrance of approximately 20 Å and is connected to the catalytic center. (b) The intramolecular cavity of naphthalene dioxygenase (PDB ID: 1o7g) with a structure similar to that of BCG3185c, identified through a query using the Dali server. These cavities are visualized using PyMOL software with the setting “set cavity_cull, 40.”

**Figure. 3.**
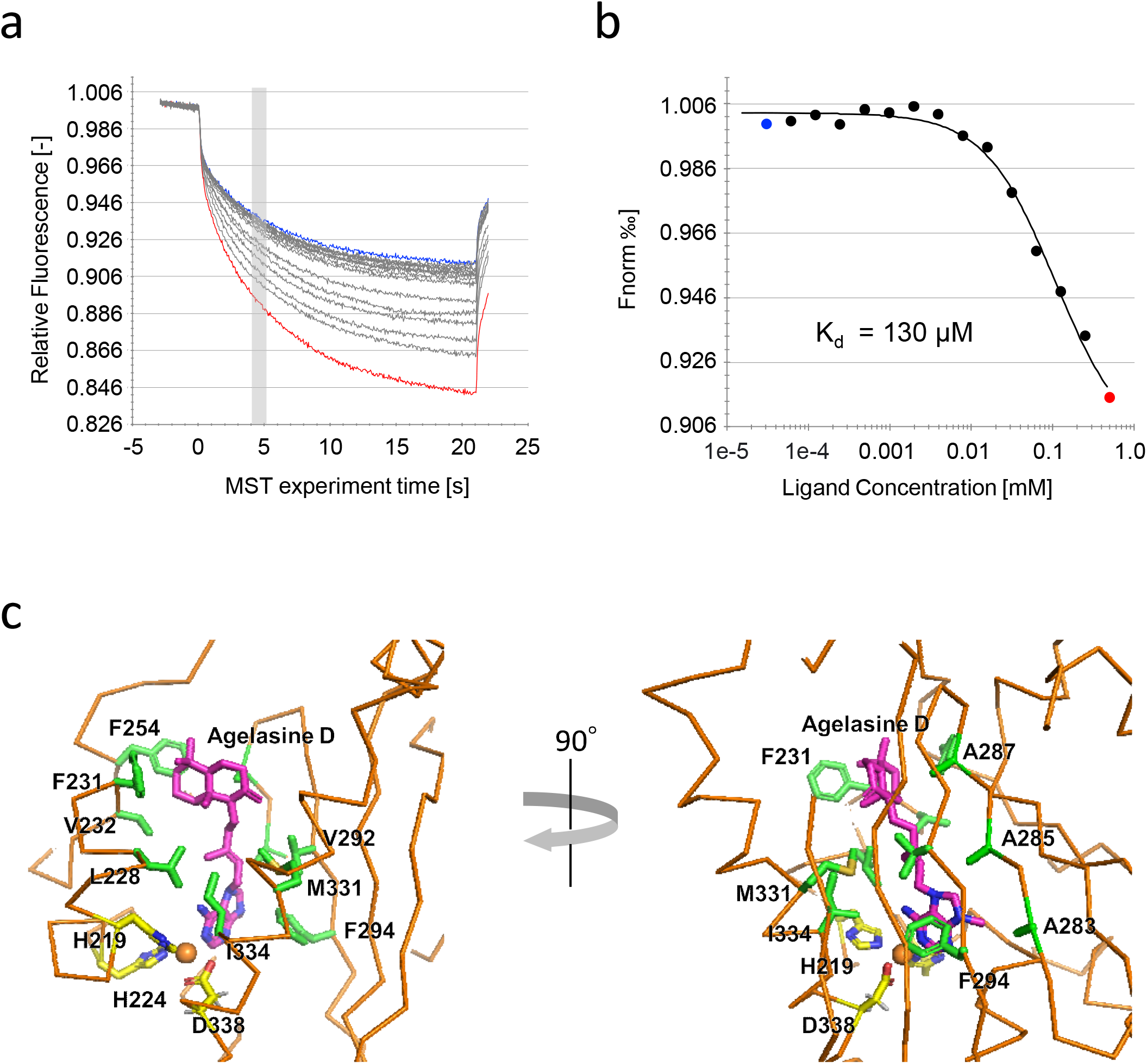
BCG3185c and agelasine D interaction analysis using microscale thermophoresis (MST) (a) The interaction of BCG3185c with agelasine D is monitored using fluorescent molecular signals attached to BCG3185c. Representative fluorescence signal traces of agelasine D titration are illustrated. The red and blue traces represent highest and lowest concentrations of agelasine D, respectively. (b) Fittings of the alteration in fluorescence signals using the dissociation constant (Kd) equation are also illustrated. The red and blue traces in Figure 3a and plots in Figure 3b represent the highest (0.5 mM) and lowest (30.6 nM) concentrations of agelasine D, respectively. The estimated Kd value for this assay is 130 μM at a BCG3185c concentration of 60 nM. (c) Binding model of agelasine D to BCG3185c through docking simulations. Multiple amino acids interact with agelasine D in its binding pocket.

## Acknowledgements

This study was supported by Grants-in-Aid from the Koyanagi-Foundation (K. Takeshita), a Grant-in-Aid for Scientific Research (C) [grant number 21K06555] (T. Yamashita) and a Grant-in-Aid for Scientific Research (B) [grant number. 21H02069] from the Japan Society for the Promotion of Science (JSPS) (M. Arai). Moreover, the Platform Project for Supporting Drug Discovery and Life Science Research (BINDS) from AMED also has supported this research in part [grant number JP19am 0101070 (support number 1328)] (T. Yamashita). We would like to thank Editage (www.editage.jp) for English language editing.

## Material & Methods

### Protein production

The BCG3185c gene was sub-cloned to the plasmid pET-47b (+) between BamHI and HindIII restriction sites, which adds an N-terminal 6xHis-tag and HRV3C recognition sequence. This plasmid was transformed into the *E. coli* BL21(DE3)RILP. The transformed cells were grown in LB media to OD_600_ = 0.8 at 37°C, and then protein expression was induced with 0.4 mM Isopropyl β-D-1-thiogalactopyranoside (IPTG) overnight at 26°C. The collected cells were sonicated with an ultrasonic system UD211 (TOMY Digital Biology co., LTD., Japan). The supernatant included recombinant BCG3185c after centrifugation (100,000g x 1hr) was purified with a nickel-affinity resin (Ni-NTA, Qiagen). The 6xHis-tag was divided from BCG3185c with digestion by HRV3C protease while dialyzing against 20 mM HepesNa, 350 mM NaCl, 0.5 mM DTT (pH 7.0). To remove 6xHis tag, HRV3C and undigested product, The protease reaction mixture was applied to Ni-NTA. The flow-through fraction containing BCG3185c was purified by size-exclusion chromatography with HiLoad 16/60 Superdex 200 prep grade (Cytiva) in 150 mM NaCl and 20 mM HEPES-Na (pH 7.0).

### X-ray crystallography

For crystallization, purified BCG3185c was concentrated to 15 mg ml ^-1^ by ultrafiltration, and crystals were obtained using hanging-drop vapor diffusion. The crystals grew from mixtures of 1 μl of protein solution and 1 μl of reservoir solution containing 4.4 M Ammonium Acetate, 0.1 M Tris-HCl (pH 8.5). All X-ray experiments were performed under a cryostream at 90 K using PEG200 as cryoprotectant. Native diffraction data was collected with X-ray wavelength of 0.90000 Å. These diffraction experiments were performed on BL26B2, BL38B1, BL44XU at SPring-8 (Harima, Japan). Finally, most high-resolution native data and each anomalous scattering data were collected at BL26B2. All diffraction data were processed and scaled with XDS^23^. All crystals belonged to space group *P*2_1_3. The structure of BCG3185c was determined determined using molecular replacement program: MOLREP^24^ and Maximum Likelihood refinement program: REFMAC5^25^. All crystallographic data are summarized in Supplementary Table 1.

### Affinity analysis of BCG3185c and Agelasine D

MicroScale Thermophoresis (MST) analysis for affinity analysis between BCG3185c and Agelasine D was performed by Monolith NT.115 (NanoTemper Technologies GmbH), following the standard assay protocols^26,27^. BCG3185c was labeled using Monolith Protein Labeling Kit RED-NHS 2nd Generation (NanoTemper Technologies Gmb) according to the protocol. BCG3185c was dissolved in PBS containing 0.02 % Tween 20 which is preventing BCG3185c aggregation and Agelasine D was dissolved in DMSO. BCG3185c was used as a binding target for the MST analysis at a concentration of 60 nM. Agelasine D, a ligand for the MST analysis, was titrated into the BCG3185c solution in a 1:1 dilution series over a concentration range (0.5 mM to 30.6 nM). BCG3185c and diluted ligand samples were mixed and loaded into a Monolith premium capillary (NanoTemper Technologies GmbH) and measured the MST fluorescence at 25°C using a MO.Control software bundled with Monolith NT.115, in which the excitation power was set as automation (96%) and the MST power was set as medium. Data were analyzed using a MO.Affinity Analysis software (version 3.0.4, NanoTemper Technologies GmbH) under standard and default MST-on time conditions (4-5 seconds: gray area in Fig. 5a). The K_d_ value was estimated by fitting the data with the equation in the software:

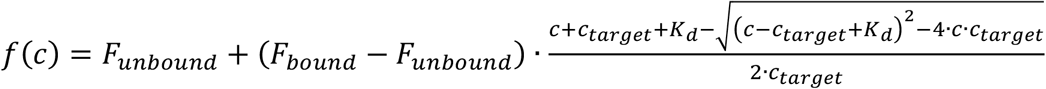

Where f(c) is the fraction bound at a given ligand concentration c; F_unbound_ is the F_norm_ signal of the target alone; F_bound_ is the F_norm_ signal of the complex; K_d_ is the dissociation constant; and C_target_ is the final concentration of target in the assay.

### Docking stimulation

Docking stimulation between BCG3185c and Agelasine D was performed by several programs in Schrödinger Suite ver. 10.3 (Schrödinger K.K., USA). Briefly, the BCG3185c structure was prepared with Prime module and Protein Preparation Wizard. Also, Agelasine D was prepared with LigPrep module in the condition pH 7.0 ± 0.5 and then docked into the prepared structure of BCG3185c using Glide module in SP mode. The result showed BCG3185c could have two Agelasine D binding sites.

## Supplementary information

**Supplementary Figure 1.**
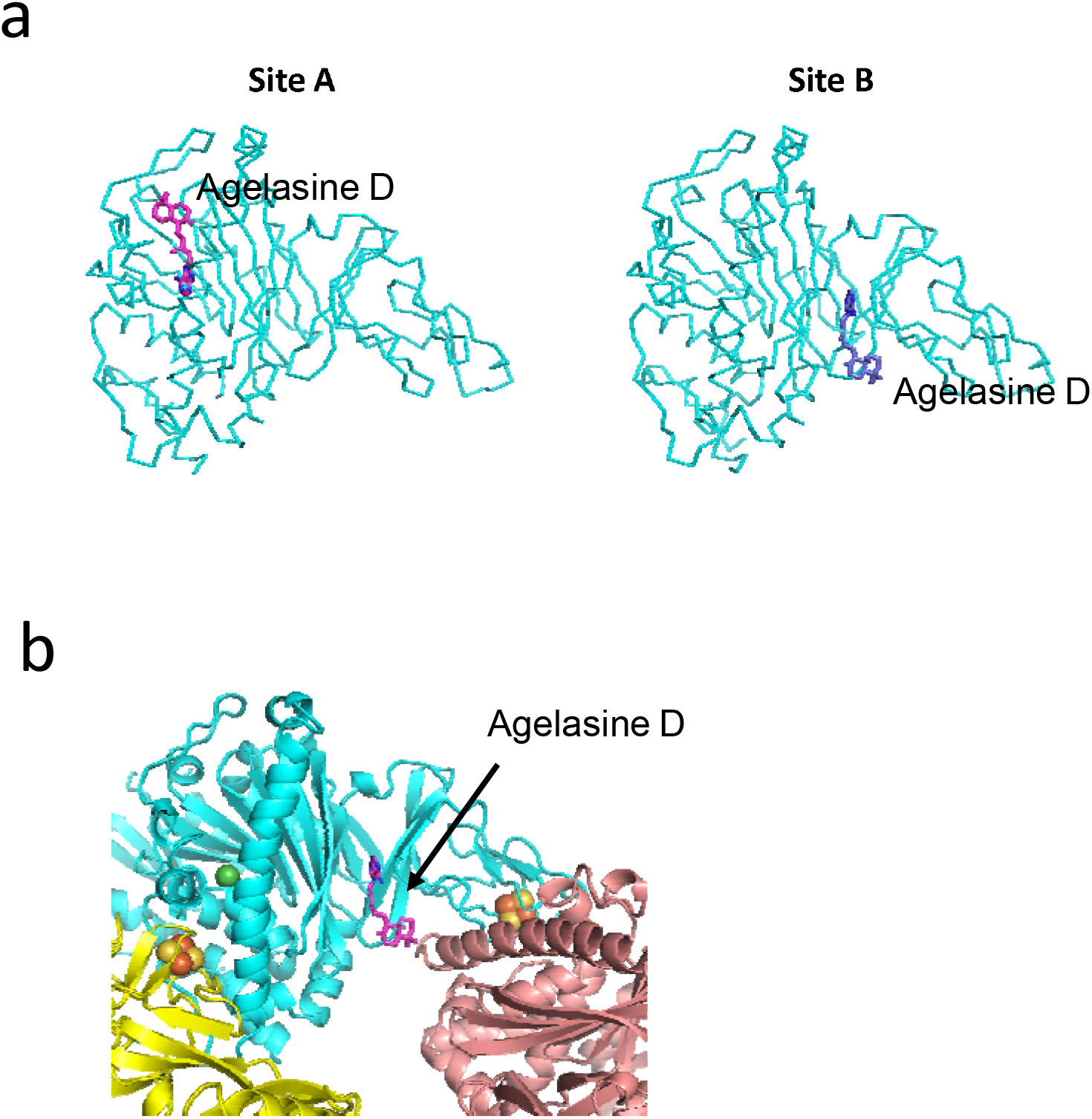
(a) Two predicted binding sites for agelasine D in BCG3185c are identified. Site A corresponds to a large hydrophobic pocket that leads to the catalytic center, whereas site B is located on the molecular surface. (b) The positioning of site B dis not disrupt the molecular interaction surface during BCG3185c trimer formation.

---

**Supplemenrary Table 1.**
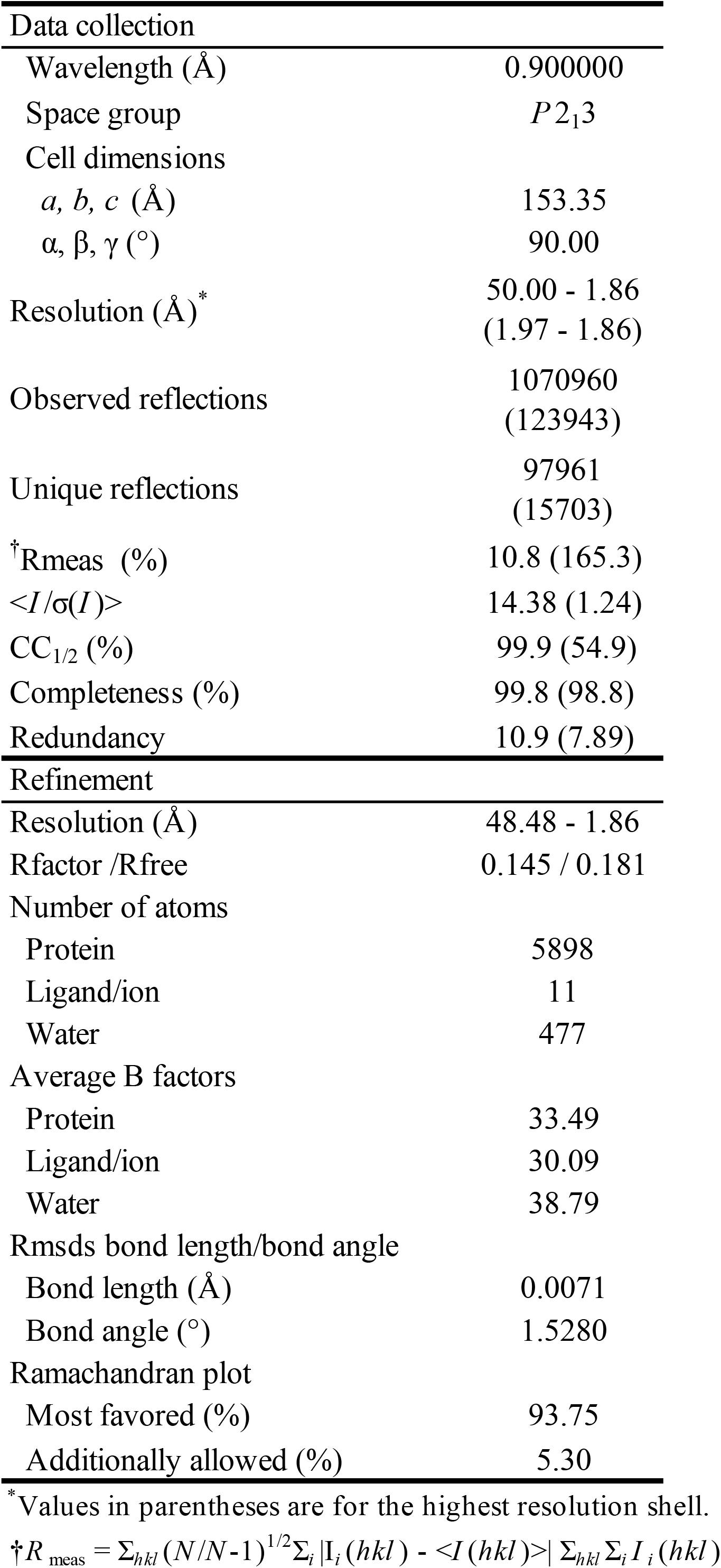
Takeshita et al..

|  |  |
| --- | --- |
| Data collection |  |
| Wavelength (Å) | 0.900000 |
| Space group | *P*2<sub>1</sub>3 |
| Cell dimensions |  |
| *a*, *b*, *c* (Å) | 153.35 |
| α, β, γ (°) | 90.00 |
| Resolution (Å) * | 50.00 - 1.86<br>(1.97 - 1.86) |
| Observed reflections | 1070960<br>(123943) |
| Unique reflections | 97961<br>(15703) |
| †R<sub>meas</sub> (%) | 10.8 (165.3) |
| <*I*/σ(*I*)> | 14.38 (1.24) |
| CC<sub>1/2</sub> (%) | 99.9 (54.9) |
| Completeness (%) | 99.8 (98.8) |
| Redundancy | 10.9 (7.89) |
| Refinement |  |
| Resolution (Å) | 48.48 - 1.86 |
| Rfactor /Rfree | 0.145 / 0.181 |
| Number of atoms |  |
| Protein | 5898 |
| Ligand/ion | 11 |
| Water | 477 |
| Average B factors |  |
| Protein | 33.49 |
| Ligand/ion | 30.09 |
| Water | 38.79 |
| Rmsds bond length/bond angle |  |
| Bond length (Å) | 0.0071 |
| Bond angle (°) | 1.5280 |
| Ramachandran plot |  |
| Most favored (%) | 93.75 |
| Additionally allowed (%) | 5.30 |

\*Values in parentheses are for the highest resolution shell.

